# Nanopore sequencing detects structural variants in cancer

**DOI:** 10.1101/028290

**Authors:** Alexis L. Norris, Rachael E. Workman, Yunfan Fan, James R. Eshleman, Winston Timp

## Abstract

Despite advances in sequencing, structural variants (SVs) remain difficult to reliably detect due to the short read length (<300bp) of 2^nd^ generation sequencing. Not only do the reads (or paired-end reads) need to straddle a breakpoint, but repetitive elements often lead to ambiguities in the alignment of short reads. We propose to use the long-reads (up to 20kb) possible with 3^rd^ generation sequencing, specifically nanopore sequencing on the MinION. Nanopore sequencing relies on a similar concept to a Coulter counter, reading the DNA sequence from the change in electrical current resulting from a DNA strand being forced through a nanometer-sized pore embedded in a membrane. Though nanopore sequencing currently has a relatively high mismatch rate that precludes base substitution and small frameshift mutation detection, its accuracy is sufficient for SV detection because of its long reads. In fact, long reads in some cases may improve SV detection efficiency.

We have tested nanopore sequencing to detect a series of well-characterized SVs, including large deletions, inversions, and translocations that inactivate the *CDKN2A/p16* and *SMAD4/DPC4* tumor suppressor genes in pancreatic cancer. Using PCR amplicon mixes, we have demonstrated that nanopore sequencing can detect large deletions, translocations and inversions at dilutions as low as 1:100, with as few as 500 reads per sample. Given the speed, small footprint, and low capital cost, nanopore sequencing could become the ideal tool for the low-level detection of cancer-associated SVs needed for molecular relapse, early detection, or therapeutic monitoring.

---

Author disclosure: Dr. Timp holds two patents (US2011/0226623 A1 and US2012/0040343 A1) which have been licensed by Oxford Nanopore Technologies.

Supported by: The Sol Goldman Pancreatic Cancer Research Center (pilot project) (JRE), SPORE P50-CA62924-13 (Kern), The Stringer Foundation (JRE), CEGS HG003323-09 (WT) and Michael Rolfe Pancreatic Cancer Foundation (JRE).

Abbreviations: Structural Variation (SV), Tumor Suppressor Gene (TSG), Pancreatic Ductal Adenocarcinoma (PDAC), Polymerase Chain Reaction (PCR)

## INTRODUCTION

Structural variants (SVs) are a hallmark of the genomic instability that underlies cancer, and include translocations, large deletions, amplifications, and inversions^1–3^. SVs are often driver alterations, with translocations and amplifications activating oncogenes and deletions and inversions inactivating tumor suppressor genes (TSGs). *CDKN2A/p16* and *SMAD4/DPC4* are two of the most commonly deleted TSGs in human cancer, and complex SVs have been found to underlie approximately half of these deletions in pancreatic ductal adenocarcinoma (PDAC)^4–6^.

The sensitive detection of tumor-specific mutations, including both small alterations such as single base substitutions and large alterations such as SVs, of circulating tumor DNA is critical for applications such as molecular relapse^7^, early detection^8^, and possibly therapeutic monitoring of cancer patients^9^. The arrival of 2^nd^ generation sequencing has provided ample opportunity to investigate small alterations, but the large SV alterations remain under-studied because of the difficulty detecting them with the short reads (<300bp) of 2^nd^ generation sequencing. Not only do the paired-end reads need to straddle a breakpoint, but repetitive elements often lead to ambiguities in the alignment. Given that repetitive regions (including centromeres, telomeres, and other repetitive elements) encompass over half (56%) of the human genome, this is a significant concern when mapping SVs^10^. The long reads (up to 20kb) generated by 3^rd^ generation DNA sequencing strategies can easily straddle these repetitive regions, allowing for unique alignment^11^.

Until recently, 3^rd^ generation sequencing was limited to PacBio, which requires a high capital investment, a large footprint, and technical expertise. These factors limit the utility of PacBio-based 3^rd^ generation sequencing in clinical testing. The new 3^rd^ generation sequencing platform, the MinION™ (Oxford Nanopore Technologies™), lacks the prohibitive factors of PacBio. The MinION instrument is the size of a large USB stick, with low (~$1k) capital cost and easy operation. Thus, nanopore sequencing on a MinION instrument may prove to be a valuable tool for clinical testing.

Nanopore sequencing, first proposed by Church et al^12^ operates via a similar principle to a Coulter counter, using a measurement of the current through a hole in a membrane to characterize sample passing through the hole. In the case of nanopore sequencing, the hole is nanometers in diameter, and the DNA molecule passing through the pore influences the current in a way which is characteristic of the local base sequence. The MinION device consists of 512 independently addressed measurement channels, each with 4 sensor wells. The software controlling the MinION selects the “best” sensor during a process called multiplexing, a process repeated several times throughout a sequencing run. Each sensor well has a semi-synthetic membrane containing a proprietary protein pore molecule. An electric field is applied across the membrane, allowing both current measurement and providing the motive force for driving the negatively charged DNA molecule through the pore. The DNA library is enriched via the tether at the membrane surface, and then diffuses along the membrane. When the DNA leader is within range of the pore, it is captured and driven through up to the motor protein. The DNA is driven through pore primarily via electric field acting on the charged phosphate backbone, with translocation velocity controlled by a proprietary motor protein coupled to the DNA molecule. The pore is large enough only for a single stranded DNA molecule; 5 bases are within the central constriction of the pore at a given time and have a significant influence on the current. After the forward DNA strand has completely run through, the hairpin is run, then the reverse or complementary DNA strand is also sequenced. The consensus of the top and bottom reads is termed a “2D read” and increases the accuracy of base calling.

Though nanopore sequencing method still has high error rate, it is rapidly improving; in our hands v7 flowcells had an average of 67.4% of the read correct, 24.2% mismatched, 7.5% insertions and 8.3% deletions^13^, but the newer v7.3 flowcells had an average of 86% correct, with 9.7% mismatch, 4.2% insertion and 4.4% deletion – a dramatic improvement (Supplemental Figure 1). Though the high error rate currently precludes their application to detecting single base substitutions (KRAS codons 12 and 13^14^) and small frameshift mutations^15,16^, the long read length easily generated with nanopore sequencing, i.e. average of 8kb reads^13^, enables easy detection of SV even in repetitive regions.

In this paper, we demonstrate the ability of nanopore sequencing to detect SVs that inactivate the *p16* and *SMAD4* TSGs in PDAC cancer cell lines. Our set of 10 SVs includes large deletions, translocations, inversions, and the complex combination of a translocation and inversion. These SVs were previously defined by SNP microarray and whole genome sequencing (WGS), and confirmed by PCR and Sanger sequencing across the junctions. We show proof-of-principle, using dilutions of PCR products containing these SVs into the corresponding wildtype amplicons and show the ability to detect these SVs at 1:100 dilutions.

## RESULTS

### Ability to detect simple and complex SVs

To demonstrate the value of long read sequencing to detect SVs, we selected a panel of 10 well-characterized SVs in the genes *CDKN2A/p16* and *SMAD4/DPC4* identified in pancreatic cancer cell lines^6^. These 10 SVs included 2 interstitial deletions, 4 translocations, 4 inversions, and 1 combination of an inversion and translocation (“TransFlip” mutations, Table 1). Wildtype (WT) sequence (intact genomic sequence; no SV) served as a control and one SV (SV01) had a technical replicate (SV07). Using Oxford Nanopore barcodes, libraries for all 12 PCR amplicons were generated and multiplexed in one flowcell run (Figure 1). The run produced a total of 3,987 2D reads from 194 of 512 channels, resulting in a 2.5Mb yield. The average read was 640bp long, full-length of our PCR products, and an average PHRED score of 11.50 (Figure 2 A-B, Supplemental Table 1). Importantly, nanopore reads have no discernible quality dependence with length, compared to the cycle dephasing commonly seen in 2^nd^ generation sequencing (Supplemental Figure 2).

**Table 1:** Details of Amplicons included in this study.

| **Amplicon ID** | **Amplicon Size Without Barcodes (bp)** | **TSG Deleted** | **SV Type** | **SV Left Breakpoint (hg19)** | **SV Right Breakpoint (hg19)** | **Expected alignment: Left (hg19)** | **Expected alignment: Center (hg19)** | **Expected alignment: Right (hg19)** |
| --- | --- | --- | --- | --- | --- | --- | --- | --- |
| SV01, SV07 | 573 | *p16* | TRANS | chr9:24,353,014 | chr22:36,338,191 | chr9:24352894-24353014(+) |  | chr22:36338191-36338601(+) |
| SV02 | 579 | *p16* | (WT) | chr9:21,970,115 | chr9:21,970,649 | chr9:21970115-21970649(-) |  |  |
| SV03 | 562 | *p16* | INV+TRANS | chr9:21,083,362 | chr9:21,083,521 | chr9:21083139-21083362(+) | chr9:21083440 - 21083521 (-) (81bp) | chr3:79387683-79387899(-) |
| SV04 | 573 | *p16* | TRANS | chr10:132,412,941 | chr9:27,096,867 | chr10:132412940-132413131(-) |  | chr9:27096866-27097203(+) |
| SV05 | 576 | *SMA D4* | ID | chr18:48,570,319 | chr18:49,191,882 | chr18:48569959-48570319(+) |  | chr18:49191882-49192052(+) |
| SV06 | 561 | *p16* | INV | chr9:24,320,470 | chr9:24,323,843 | chr9:24323843-24324156(-) |  | chr9:24320470-24320672(+) |
| SV08 | 559 | *p16* | INV | chr9:25,968,399 | chr9:25,969,868 | chr9:25968120-25968399(+) |  | chr9:25969634-25969868(-) |
| SV09 | 581 | *p16* | INV | chr9:25,969,504 | chr9:25,972,326 | chr9:25969502-25969820(-) |  | chr9:25972324-25972543(+) |
| SV10 | 584 | *p16* | INV+TRANS | chr9:21,326,884 | chr7:140,023,555 | chr9:<br>21326735-21326867(+) | chr9:<br>21326884<br>-<br>21326931<br>(-) (47bp) | chr7:<br>140023553-140023913(+) |
| SV11 | 578 | *SMA*<br>*D4* | TRANS | chr6:124,911,349 | chr18:53,465,051 | chr6:124911<br>349-124911707(-<br>) |  | chr18:5346504<br>9-53465220(+) |
| SV12 | 573 | *SMA*<br>*D4* | ID | chr18:48,434,141 | chr18:49,851,882 | chr18:48433<br>731-48434141(+) |  | chr18:4985188<br>2-49852004(+) |
| *Tumor Suppressor Gene (TSG), Structural Variant (SV), Translocation (TRANS), Wild-type Control (WT), Inversion (INV), Interstitial Deletion (ID)* |  |  |  |  |  |  |  |  |

**Figure 1:**
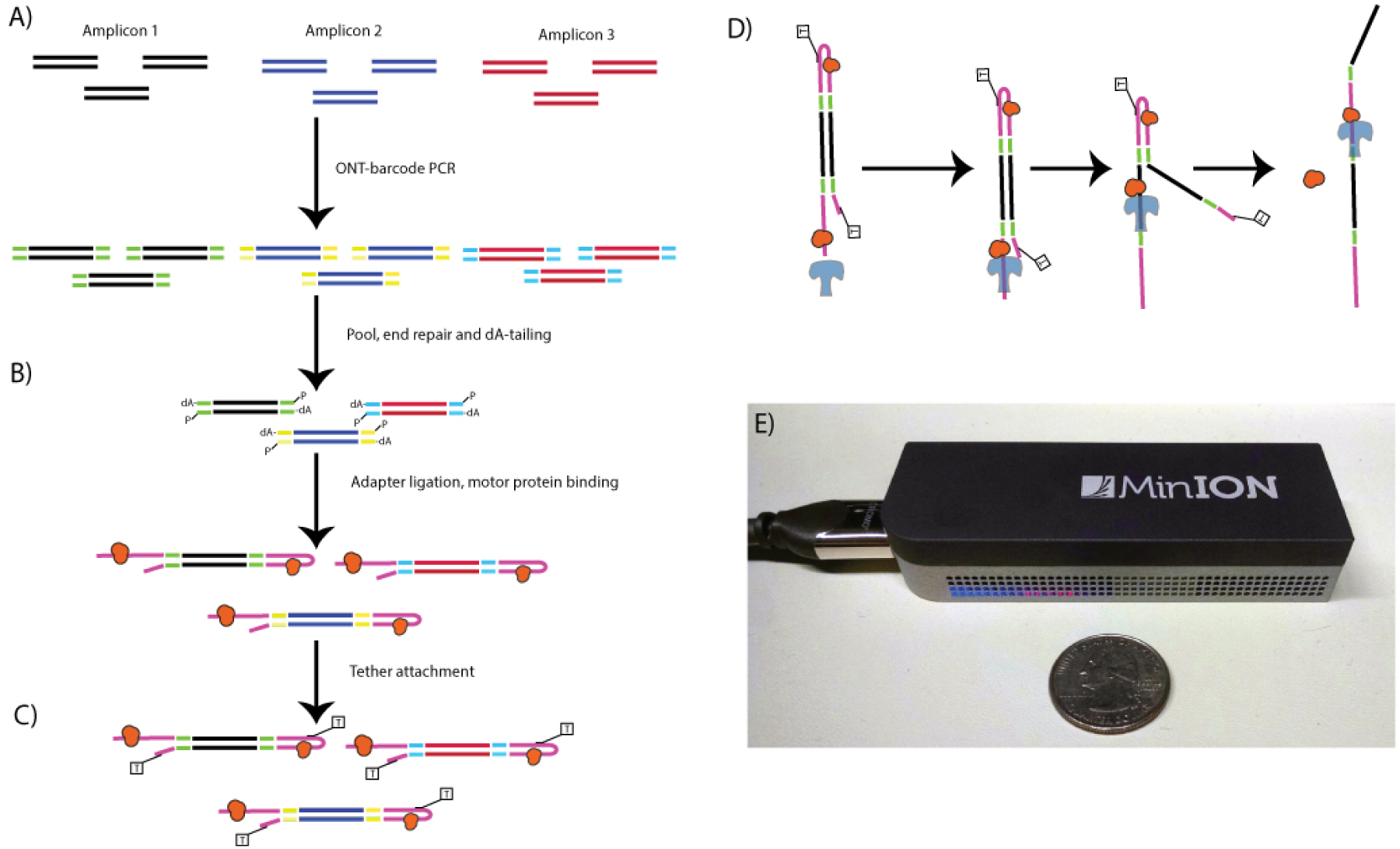
Nanopore Library Prep Workflow. Oxford Nanopore barcodes were incorporated into amplicons by PCR-individually for each SV-then resultant reactions were pooled (A). After NEB End Repair and dA-tailing modules (B), hairpin and leader adapters were ligated on, each containing a motor protein. Only the hairpin protein contained a his-tag, which was used to enrich for molecules containing a leader adapter and his-tag (his-tag selection step not shown). Tether attachment (C) allowed for direct attachment of the molecules to the flow cell membrane. Within the MinION flowcell (D), DNA molecules are pulled through a protein pore (blue), with motor protein (orange) affecting speed of DNA translocation through the pore. One side of the DNA molecule is read, then the hairpin, then the second side. Both reads were aligned to produce a 2D consensus read.

**Figure 2:**
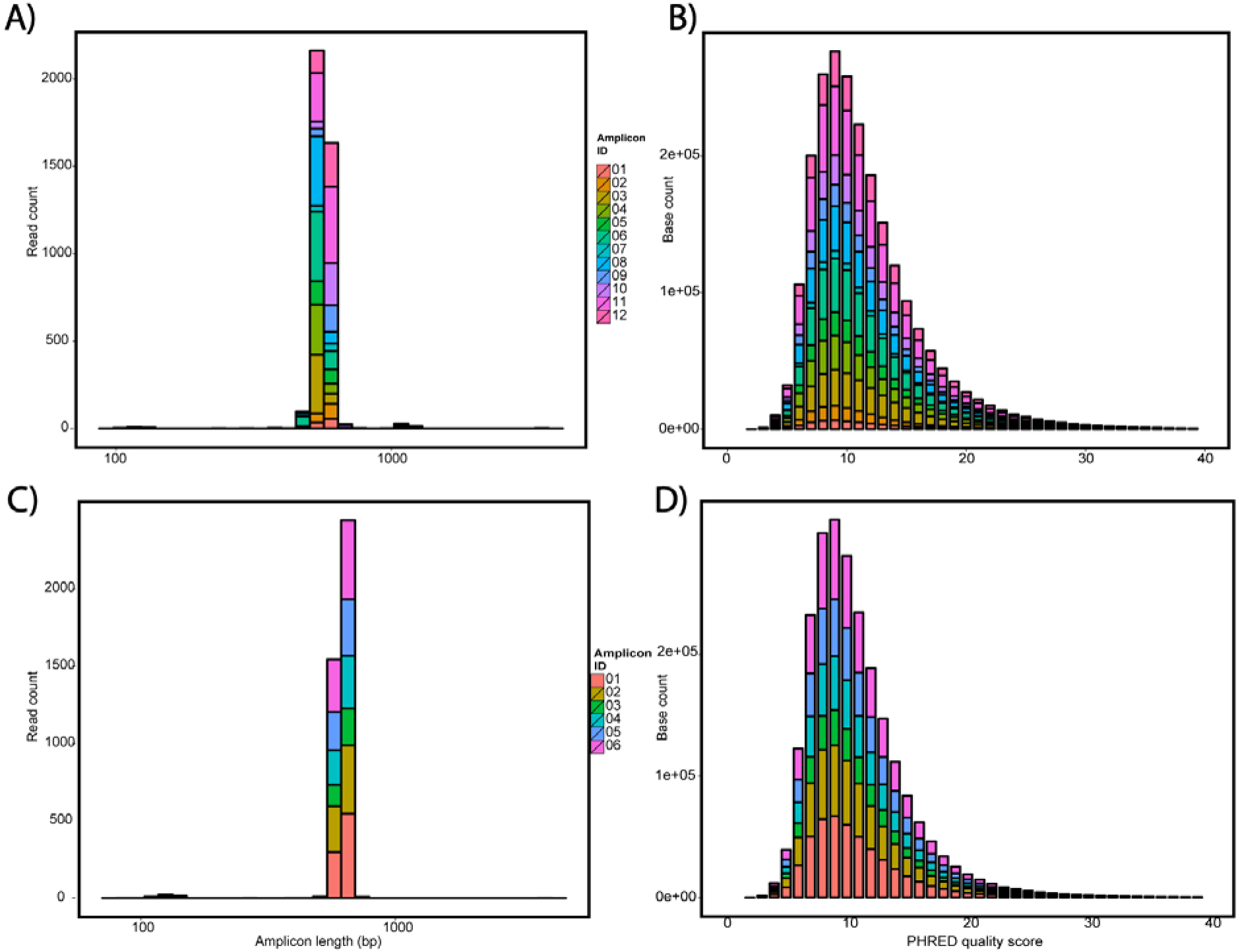
Nanopore sequencing QC data. QC of Flow cell 1 A) length and B) PHRED quality histograms of each of the barcodes as a stacked bar graph. Average length of 570 bp and PHRED score of 11.5. QC of flow cell 2 C) length and D) PHRED quality histograms. Average length of 573 bp and PHRED score of 10.9

All SV amplicons (12/12) mapped to their expected region(s) of hg19 (Table 2, Supplemental Figures 3-6), with overall mapping percentage of 99.6% and 79% of aligned reads with correctly matched bases (Supplemental Table 2). Importantly, the representation of the SV amplicons seems independent of the complexity of their SV (intact genomic sequence (SV02) represented 3.5% of aligned reads, and a complex combination of a deletion, inversion, and translocation (SV10) represented 5.1% of aligned reads (Table 2). The technical replicates (SV01, SV07) had comparable results, as expected. However, some amplicons had a surprisingly low percentage of properly aligned SV structures, specifically of note is SV03 with only 16.5% correctly aligned. In this case only 68 (16.5%) reads had the full alignment of left, center and right alignment. However, 313 reads (76.0%) had the left and right alignment, and 12 further reads (2.9%) had left and center alignment.

**Table 2:** All SVs are detected by Nanopore multiplex (1:12) experiment [Exp1].

| Amplicon ID | SV Type | 2D reads | % total reads per barcode | 2D reads aligned to hg19 (%) | Reads properly aligned* (%) | Off-target Reads | Lumpy break-points | Top Lumpy Breakpoint |
| --- | --- | --- | --- | --- | --- | --- | --- | --- |
| SV01 | TRANS | 100 | 2.5% | 91 (91.0%) | 77 (77.0%) | 6 (6.0%) | 1 | chr9:24353014 /chr22:36338191 (58) |
| SV02 | n/a (WT) | 140 | 3.5% | 139 (99.3%) | 115 (82.1%) | 24 (17.1%) | 0 | None |
| SV03 | INV+TRANS | 412 | 10.3% | 412 (100.0%) | 68 (16.5%) | 1 (0.2%) | 8 | chr3:79387939 /chr9:21083384 (132) |
| SV04 | TRANS | 356 | 8.9% | 356 (100.0%) | 303 (85.1%) | 6 (1.7%) | 3 | chr9:27096843 /chr10:132412942 (183) |
| SV05 | ID | 225 | 5.6% | 224 (99.6%) | 198 (88.0%) | 7 (3.1%) | 2 | chr18:48570319 (154) |
| SV06 | INV | 567 | 14.2% | 567 (100.0%) | 549 (96.8%) | 7 (1.2%) | 4 | chr9:24320456 /chr9:24323864 (120) |
| SV07 | TRANS | 80 | 2.0% | 78 (97.5%) | 70 (87.5%) | 0 (0.0%) | 2 | chr9:24353014 /chr22:36338191 (52) |
| SV08 | INV | 487 | 12.2% | 487 (100.0%) | 449 (92.2%) | 3 (0.6%) | 4 | chr9:25968397 /chr9:25969868 (384) |
| SV09 | INV | 202 | 5.1% | 202 (100.0%) | 190 (94.1%) | 5 (2.5%) | 2 | chr9:25969501 /chr9:25972324 (172) |
| SV10 | INV+TRANS | 306 | 7.7% | 301 (98.4%) | 254 (83.0%) | 1 (0.3%) | 5 | chr7:140023522 /chr9:21326914 (167) |
| SV11 | TRANS | 725 | 18.2% | 725 (100.0%) | 471 (65.0%) | 3 (0.4%) | 3 | chr6:124911349 /chr18:53465049 (362) |
| SV12 | ID | 387 | 9.7% | 387 (100.0%) | 274 (70.8%) | 3 (0.8%) | 2 | chr18:48434141 (115) |
| **Average** |  | **332** | **8.3%** | **331 (98.8%)** | **252 (78.2%)** | **6 (2.8%)** |  |  |
| *To be considered properly aligned, a read must align to all expected regions (eg. Left sequence, Center sequence, and Right sequence, from Table 1) |  |  |  |  |  |  |  |  |

### Ability to detect low frequency SVs

We next wanted to determine the sensitivity of nanopore based SV detection to low frequency or rare events, to simulate a clinical scenario. To this end, we performed a 1:100 dilution of 6 SV amplicons in a background of intact *p16* genomic sequence (SV02, Table 3). These 6 amplicons included 2 simple interstitial deletions, 2 translocations, 1 inversion, and 1 complex combination of an inversion and translocation. The run produced a total of 4,058 2D reads from 270 of 512 channels, for a total yield of 2.6Mb, with an average read length of 650bp and an average PHRED score of 10.9 (Figure 2C-D, Supplemental Tables 3-4). All 6 SV amplicons were represented in the alignment (range 9-21%) and aligned to the expected regions of hg19 (Table 3). Remarkably, even with only 378 2D reads in the case of SV03, the SV was detected, with 10 of the 378 reads supporting a t(9;22).

**Table 3:** Results of low frequency serial dilutions of SVs 1:100 into wildtype [Exp2].

| Amplicon ID | SV Type | Dilution into WT | # 2D reads | % per barcode | Reads aligned to hg19 | Aligned reads mapped to WT | Aligned reads mapped to SV* | # Off-target Reads | Lumpy break-points | Top Lumpy Breakpoint |
| --- | --- | --- | --- | --- | --- | --- | --- | --- | --- | --- |
| SV01 | TRANS | 1:100 | 867 | 21.37% | 851<br>(98.2%) | 838<br>(96.7%) | 11<br>(1.3%) | 3<br>(0.3%) | 1 | chr9:24353014/<br>chr22:36338197 (7) |
| SV03 | INV+<br>TRANS | 1:100 | 760 | 18.73% | 741<br>(97.5%) | 685<br>(90.1%) | 7 (0.9%) | 50<br>(6.6%) | 3 | chr3:79387933/<br>chr9:21083384 (13) |
| SV04 | TRANS | 1:100 | 378 | 9.31% | 377<br>(99.7%) | 367<br>(97.1%) | 10<br>(2.6%) | 1<br>(0.3%) | 1 | chr9:27096848/<br>chr10:132412942 (8) |
| SV05 | ID | 1:100 | 577 | 14.22% | 571<br>(99.0%) | 538<br>(93.2%) | 31<br>(5.4%) | 3<br>(0.5%) | 1 | chr18:48570319 (25) |
| SV09 | INV | 1:100 | 621 | 15.30% | 617<br>(99.4%) | 601<br>(96.8%) | 16<br>(2.6%) | 1<br>(0.2%) | 1 | chr9:25969504/<br>chr9:25972324 (14) |
| SV12 | ID | 1:100 | 855 | 21.07% | 849<br>(99.3%) | 810<br>(94.7%) | 26<br>(3.0%) | 14<br>(1.6%) | 1 | chr18:48434141 (11) |
| **Mean** |  |  | **676** | **16.67%** | 668<br>(98.8%) | 640<br>(94.8%) | 17<br>(2.6%) | 12<br>(1.6%) |  |  |
| *To be considered properly aligned, a read must align to all expected regions (eg. Left sequence, Center sequence, and Right sequence, from Table 1) |  |  |  |  |  |  |  |  |  |  |

### SV breakpoint location detection

Finally, to determine the accuracy of breakpoint location detection with this new sequencing methodology, we first employed LUMPY^17^, an established tool for breakpoint detection for both discordant paired end short-read sequencing and long, split read alignments. Using the alignment files generated from BWA above, we extracted the split reads and fed the resulting BAM file into LUMPY. The results are included in Tables 2 and 3.

For some of our samples, LUMPY detected the correct breakpoint, and only one breakpoint, (SV01), or detected no breakpoint in the WT sample (SV02), but in general the breakpoints it detected, though correct in type, lacked precision. In the duplicate sample of SV01, SV07, the same correct breakpoint was detected. Not as many pieces of evidence (as decided by LUMPY) support this breakpoint as when simply examining coverage, because LUMPY has strict map quality filters which remove some of the reads from consideration. In many cases LUMPY detects many breakpoints at slightly shifted conditions – to a max of 8 breakpoints detected in SV03. The breakpoint with the plurarity of reads accepted by LUMPY isrepresented in Tables 2 and 3.

We examined the alignment more carefully to determine the cause of these artifacts in breakpoint location detection. Supplemental Figures 4-6 give a hint as to the problem – a careful examination notes that the reads frequently align past the breakpoint, but most of the bases are mismatched in these locations. With bwa bound by the promiscuous settings used for nanopore sequencing alignment, it continues the alignment past the breakpoint. We summarized these findings in Table 4. Though in many cases the alignment termini are set to the breakpoints correctly, for SV03 in particular this happens a minority of the time. When the alignment slips past the downstream boundary of the left fragment, there is no longer sufficient sequence for the center fragment to align.

**Table 4:** Alignment termini position error

| **Amplicon ID** | **Overlapping alignments** | **Correct Upstream Termini (%)** | **Mean Upstream Error ± SD** | **Correct Downstream Termini (%)** | **Mean Downstream Error ± SD** |
| --- | --- | --- | --- | --- | --- |
| SV01, SV07 (L) | 77 | 3 (3.9%) | 1.8 ± 5.4 | 53 (68.8%) | 5.3 ± 14.5 |
| SV01, SV07 (R) | 85 | 4 (4.7%) | 1.1 ± 5.5 | 1 (1.2%) | -1.3 ± 17.3 |
| SV02 | 116 | 7 (6.0%) | 0.1 ± 5.2 | 13 (11.2%) | -4.3 ± 10.1 |
| SV03 (L) | 407 | 6 (1.5%) | 4.9 ± 8.3 | 16 (3.9%) | 20.8 ± 13.7 |
| SV03 (C) | 83 | 1 (1.2%) | 10.8 ± 18.4 | 58 (69.9%) | 4.7 ± 18.7 |
| SV03 (R) | 391 | 51 (13.0%) | 1.8 ± 17.2 | 3 (0.8%) | 39.1 ± 24.2 |
| SV04 (L) | 317 | 23 (7.3%) | 11.0 ± 19.3 | 1 (0.3%) | 7.3 ± 7.0 |
| SV04 (R) | 349 | 2 (0.6%) | 19.0 ± 23.3 | 196 (56.2%) | -5.7 ± 14.2 |
| SV05 (L) | 225 | 4 (1.8%) | 5.7 ± 8.5 | 107 (47.6%) | 2.9 ± 14.2 |
| SV05 (R) | 206 | 1 (0.5%) | 7.3 ± 9.8 | 62 (30.1%) | -3.8 ± 9.7 |
| SV06 (L) | 565 | 0 (0.0%) | -20.9 ± 5.2 | 229 (40.5%) | -1.0 ± 4.5 |
| SV06 (R) | 555 | 4 (0.7%) | 5.6 ± 13.6 | 297 (53.5%) | -1.6 ± 7.8 |
| SV08 (L) | 70 | 6 (8.6%) | 2.7 ± 5.4 | 45 (64.3%) | 2.9 ± 14.7 |
| SV08 (R) | 78 | 3 (3.8%) | 4.4 ± 14.3 | 1 (1.3%) | -1.1 ± 7.9 |
| SV09 (L) | 476 | 125 (26.3%) | 4.4 ± 9.9 | 116 (24.4%) | -2.1 ± 6.6 |
| SV09 (R) | 464 | 37 (8.0%) | 0.0 ± 11.0 | 276 (59.5%) | 7.0 ± 28.4 |
| SV10 (L) | 271 | 32 (11.8%) | -0.5 ± 4.7 | 0 (0.0%) | 49.9 ± 19.4 |
| SV10 (C) | 260 | 0 (0.0%) | 144.0 ± 26.5 | 0 (0.0%) | -9.9 ± 14.7 |
| SV10 (R) | 302 | 0 (0.0%) | 28.8 ± 24.4 | 8 (2.6%) | 15.7 ± 27.6 |
| SV11 (L) | 725 | 4 (0.6%) | 34.7 ± 51.5 | 91 (12.6%) | -5.7 ± 7.9 |
| SV11 (R) | 472 | 8 (1.7%) | -0.1 ± 8.0 | 75 (15.9%) | 2.4 ± 8.6 |
| SV12 (L) | 386 | 72 (18.7%) | 5.2 ± 6.9 | 151 (39.1%) | 4.1 ± 11.2 |
| SV12 (R) | 278 | 11 (4.0%) | -0.6 ± 7.9 | 6 (2.2%) | 9.9 ± 9.9 |

## DISCUSSION

The accurate and timely detection of tumor-associated alterations, including SVs, is important for patient management, from early detection to monitoring for molecular relapse, as well as determining or predicting chemoresponse. Detection of all tumor-associated alterations is complicated by the low tumor cellularity often present in tumor samples and biopsies due to contaminating normal cells. Tumor-associated SVs are additionally complicated when they occur in repetitive regions, which account for over half of the human genome. The ability of long-read 3^rd^ generation sequencing methods, such as nanopore, to read through repetitive regions could make it an ideal tool for detecting tumor-associated SVs.

This work serves as proof-of-principle, showing the ability of nanopore sequencing to correctly and reliably detect SV with only hundreds, instead of millions of reads. Furthermore, we have demonstrated the feasibility of the MinION for the detection of well-characterized patient-specific SV rearrangements using *in vitro* mixtures of PCR amplicons at 1:100 dilutions in wildtype sequence. The 4 types of SVs assessed in this study include simple interstitial deletions, translocations, inversions, and the complex combination of a translocation and an inversion. This is accomplished despite the error rate (at the single-base level) of this emerging technology, because the read length is relatively long, the human genome is known, and this level of accuracy is sufficient to correctly map hundreds to thousands of bases, even if they contain multiple point mutation errors. Precision of breakpoint location is still limited, but this can be solved bioinformatically, via alignment parameter optimization or breakpoint detection tailored to the idiosyncrasies of nanopore sequencing data.

The primary advantages for nanopore sequencing over 2^nd^ generation sequencing methods for detection of SV are its (1) ability to sequence through repetitive regions, (2) speed, and (3) low cost and availability.

First, the long-read nature (up to 20kb) of nanopore sequencing allows reading through repetitive regions. Even with long mate-pair sequencing at deep coverage, 2^nd^ generation methods' short-read sequences prohibit accurate and efficient mapping of repetitive regions, which often house SVs. Previous work has demonstrated that long-read sequencing on its own has been able to detect novel SVs; 10% of the ~30,000 SVs detecting in a single individuals somatic genome were detected only via long-read PacBio sequencing^18^.

Second, the speed of real-time nanopore sequencing offers results in minutes, allowing for rapid diagnosis and treatment. To have 99% confidence of a variant at 1:100 in the sample, we need ~450X coverage over the region of interest^19^. In nanopore, each of the 512 channels can generate a read separately, with each read completed and analyzable in minutes. From the two sequencing runs we performed in this paper, generating 450 reads required 15 minutes and 33 minutes respectively. In contrast, 2^nd^ generation sequencing generates millions of reads simultaneously, but the reads are only complete after hours or days, meaning that any analysis has to wait for completion. For example, the fastest Illumina 2^nd^ generation sequencing run requires 4 hrs to obtain 1x36 bp 12 M reads (MiSeq v2); and such short reads would prove challenging for SV detection. In both cases we are omitting the library preparation time, but these times are largely equivalent.

Third, the low cost (approximately $1k per device) and small size (USB stick) of the MinION nanopore sequencing instrument offer accessibility to testing in nearly any setting. In contrast, the instrumentation for 2^nd^ generation methods require a substantial upfront investment (>$100k) and sufficient lab space for their large footprint, which are prohibitive to many research and clinical labs.

There are currently two limitations that restrict the utility of nanopore sequencing: (1) a relatively high mismatch and indel error rate and (2) limited yield (on the scale of Megabases or Gigabases), but both of these factors continue to improve. In our hands, error rate per read decreased from 32% to 14% over a 6 month period (Supplemental Figure 1B). Better tools for corrected basecalling^20^, alignment^21,22^ and assembly tools^23^ have already been generated by the community. While still insufficient for whole-genome sequencing, the MinION yield has been increasing, and yields per flow cell by other groups have reached nearly 2Gb, with substantially greater improvements (10 fold) in yield expected before the end of 2015. Additionally, the throughput of nanopore sequencing should increase with the release of the PromethION, GridION, and subsequent systems from Oxford Nanopore.

Here we have shown the ability and reliability of nanopore sequencing, a 3^rd^ generation sequencing method, to detect well-characterized SVs, and at low-levels that simulate that seen in the clinical testing. Importantly, the SV sequences were represented equally well in the alignments of nanopore sequencing data - from simple (interstitial deletions) to complex (inversions and translocations) SVs. Further development is needed on bioinformatics tools which can precisely align to and detect breakpoint locations. It will be critically important to demonstrate the ability to detect SVs from cancer/normal cell titrations of genomic DNA, as well as plasma from pre- and post-resection patients. Ongoing studies involve further dilution experiments and detection of novel (unknown) SVs directly from patient samples.

## MATERIALS AND METHODS

### Identification of SVs

Genomic DNA was extracted from previously described PDAC cancer cell lines using QIAamp DNA mini kit (Qiagen), per manufacturer’s instruction^24^. Structural variants associated with *p16* and *SMAD4* deletions were identified by high density SNP microarray and WGS, and confirmed by PCR amplifying across the novel DNA:DNA junction and bidirectional Sanger sequencing^6^. Primers were designed upstream and downstream of 10 *p16* and *SMAD4* deletions associated with different SVs (Table 1), as well as *p16* wildtype sequence, to produce amplicons of 550-600 basepairs^25^. We also included a technical replicate in our design to control for technical variation (SV01 and SV07). Residual nucleotides and oligonucleotides were removed using QIAquick PCR purification kit (Qiagen), per manufacturer’s instructions. PCR specificity was verified by gel electrophoresis and quantified by Qubit DNA double-stranded high sensitivity assay.

### Library Preparation

Barcodes were added to the PCR amplicons with Oxford Nanopore primers complementary to the tail sequence with a sample specific barcode (Barcode Developer Kit I) using Long Range PCR kits (NEB) (Figure 1). This allowed for multiplexing of up to 12 samples on a single flow cell. Barcoded PCR libraries were quantified with Qubit dsDNA HS Assay kit (Life Technologies), normalized, and pooled to a final amount of 1 μg. For sequencing, the libraries were end-repaired and dA-tailed using NEB DNA Ultra modules, followed by the ligation of hairpin and Oxford Nanopore-specific leader adapters using Genomic DNA Sequencing Kit MAP-004 (Oxford Nanopore). A motor protein was bound to both the leader and hairpin adapters, and serves to ratchet each molecule through the nanopore one base at a time. Enrichment for molecules containing hairpin adapters and bound motor protein was performed using His-Tag Dynabeads^®^ (Life Technologies).

### Flowcell Runs

For the first flowcell run, the 12 amplicons were multiplexed together at equal concentrations. For the second flowcell run, *in vitro* dilutions were performed to assess the ability to detect low-level SVs, to simulate clinical samples. Specifically, the following *p16-* and *SMAD4*-associated SVs were diluted at 1:100 in wildtype *p16* sequence (SV02): an inversion (SV09), an inversion with translocation (SV03), translocations (SV01 and SV04), and simple interstitial deletions (SV05 and SV12). These dilutions were barcoded and multiplexed together at equal concentrations.

### Oxford Nanopore MinION^TM^ Sequencing and Basecalling

The MinION Flow Cell (R7.3 chemistry) was run for 48 hours on MinKNOW software (v0.49.3.7), producing thousands of fast5 files, each file corresponding to a molecule read by the sequencer. Cloud-based basecalling software (Metrichor^TM^, v2.29.1, Oxford Nanopore) was used to convert electrical event data from MinKNOW into basecalled files. Three basecalled reads were produced: a “1D template” and “1D complement”, and “2D read”. The 2D read is the consensus sequence between the template and complement reads, and a basic quality filter is applied to keep only 2D reads with a ratio of template bases to complement bases between 0.5 and 2.

Nanopore basecalling is performed by Metrichor using a hidden Markov model, similar to the process described in a simulated data set previously^26^. Briefly, each pentamer generates a specific current which, although difficult to distinguish uniquely, combined with the controlled translocation rate, allows for basecalling the best full sequence.

### Alignment and SV calling of Reads

Using only 2D nanopore reads which passed the quality filter, we de-mulitplexed and extracted fastq data with custom code in python (https://github.com/timp0/timp_nanoporesv). We then aligned the nanopore long reads against the hg19 reference genome using BWA-MEM, with the – x ont2d option set for nanopore specific alignment parameters^22^. A custom python script to extract split read alignments and calculate error in alignment location is also included in the online git repository.

## Acknowledgements

We thank Oxford Nanopore for outstanding technical support. We thank Aaron Quinlan and Ryan Layer for helpful discussions.

## Supplemental Figures

**Supplemental Figure 1:**
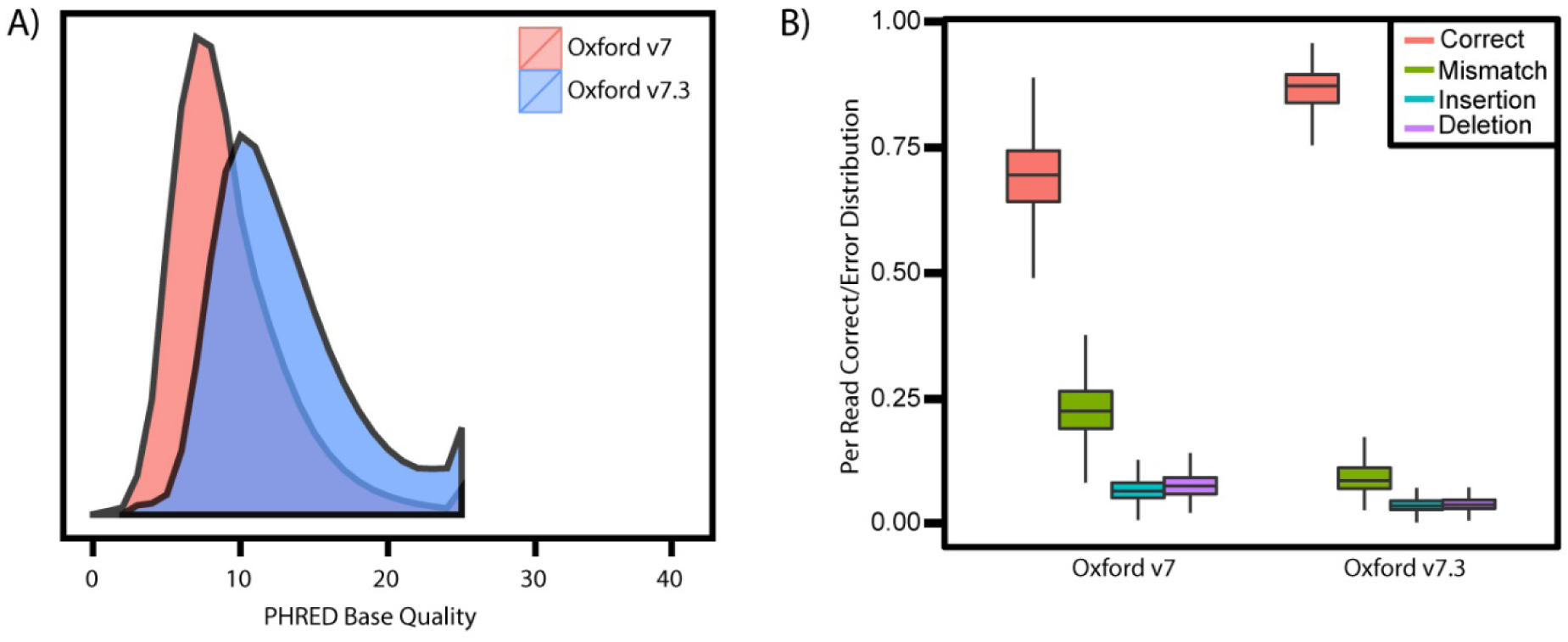
Improvement of Oxford Nanopore data quality from v7 to v7.3 on site. A) Density plot of PHRED density from FASTQ files generated from v7 and v7.3 sequencing runs. Median PHRED quality score for v7 flowcells was XX, while median PHRED score for v7.3 was YY B) Comparison of distributions of correct, mismatch, insertion and deletion events as a fraction of total read for v7 and v7.3 nanopore data. v7 flowcells had an average of 67.4% of the read correct, 24.2% mismatched, 7.5% insertions and 8.3% deletions, with v7.3 flowcells having an average of 86% correct, with 9.7% mismatch, 4.2% insertion and 4.4% deletion.

**Supplemental Figure 2:**
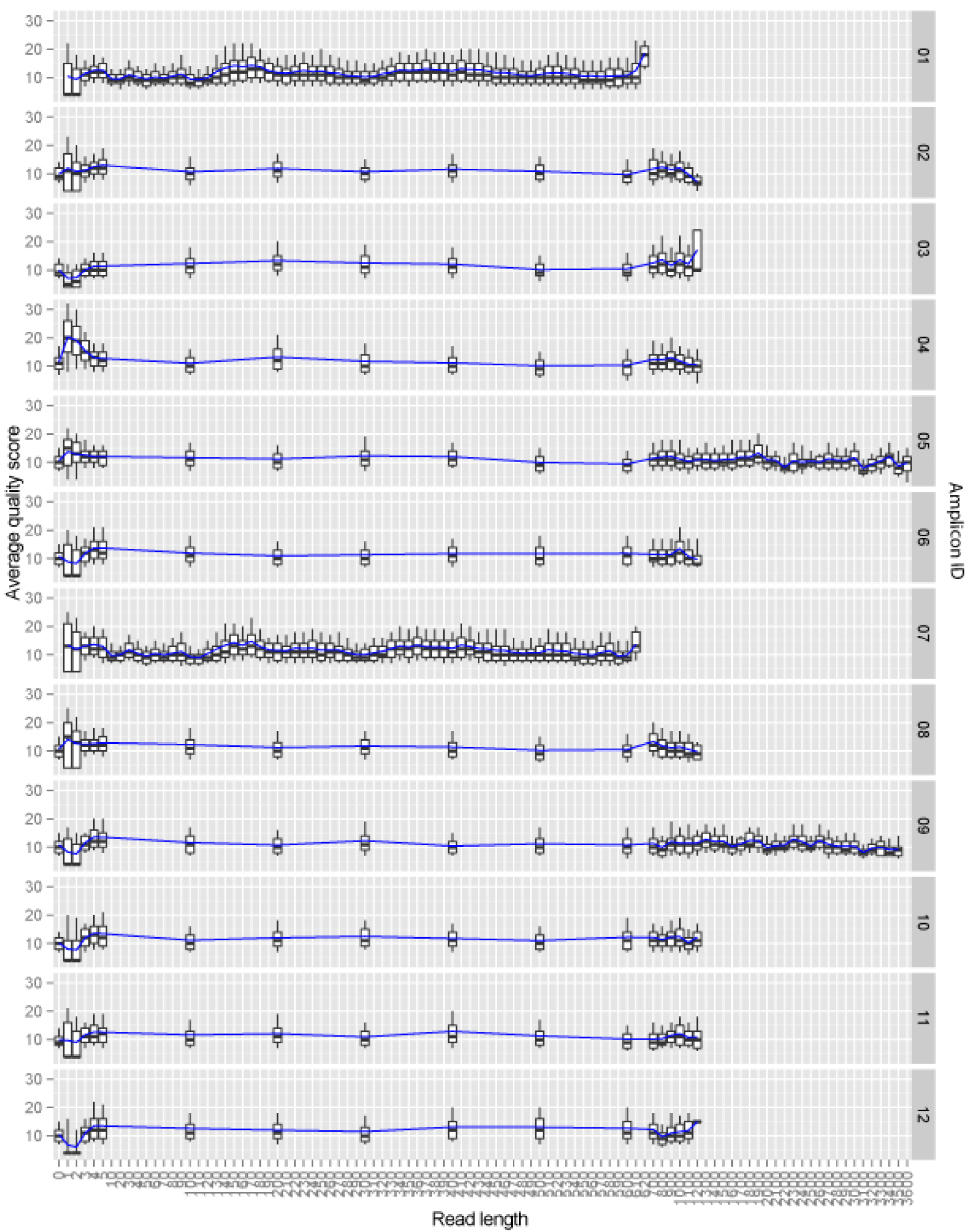
Quality score from nanopore reads versus position along read for the 12 amplicons multiplexed in the first sample run. No discernible drop in quality versus read length is present.

**Supplemental Figure 3:**
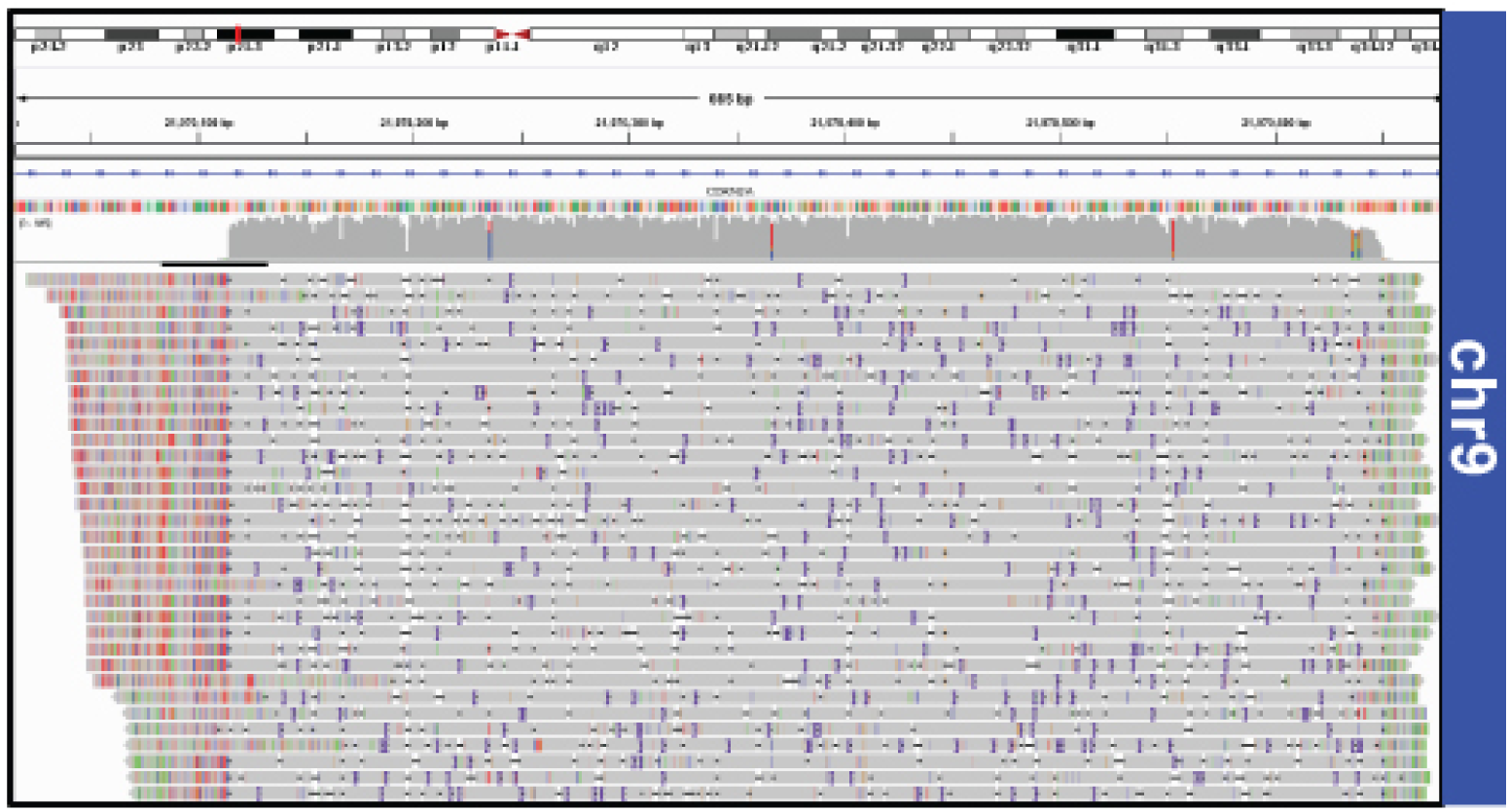
IGV screenshot of WT (SV02) alignment.

**Supplemental Figure 4:**
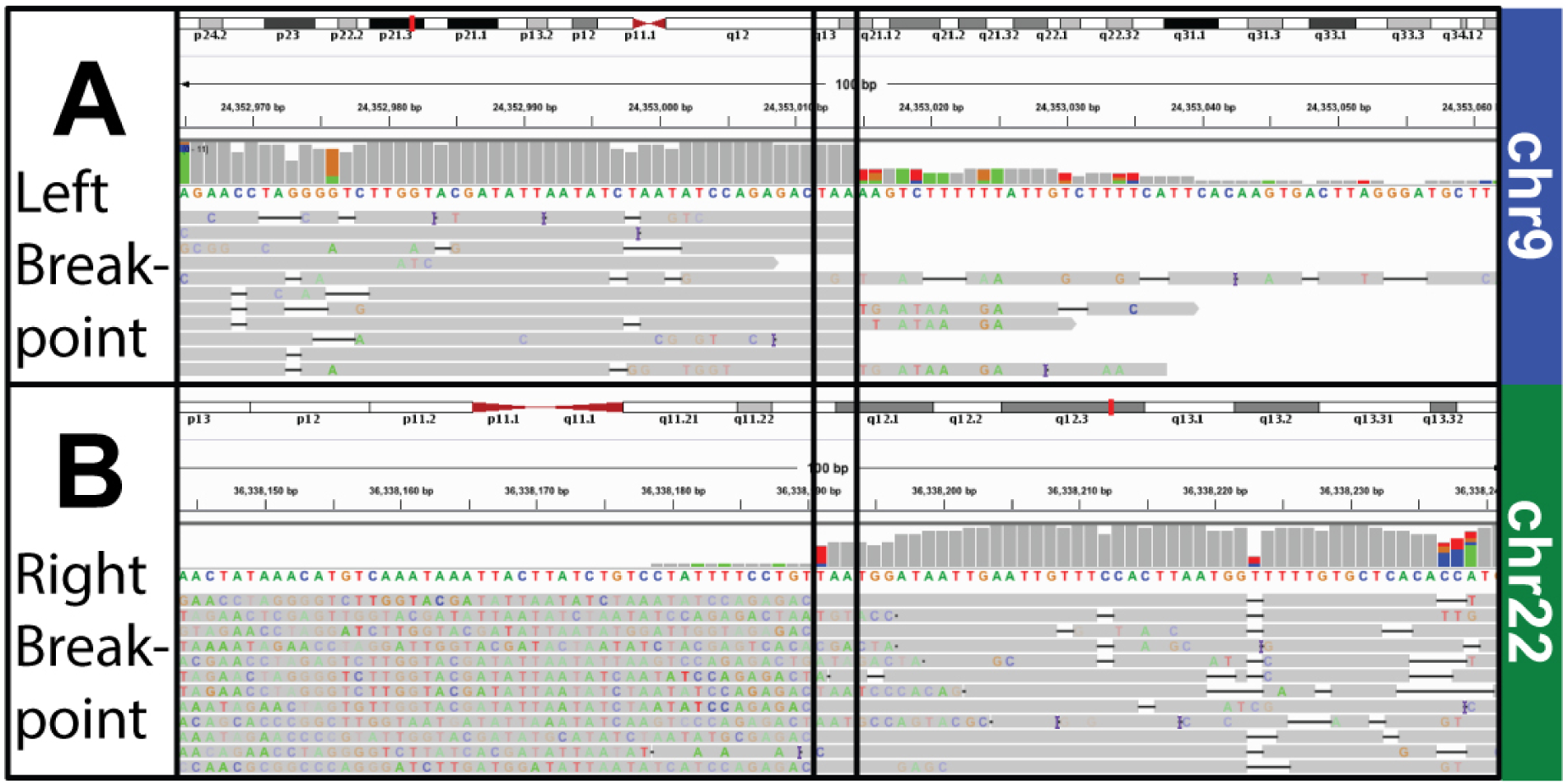
IGV Screenshot of Translocation (SV01) alignment. A) Shows the alignment to the area in chr9 and B) the alignment to the area in chr22. Note the erroneous extension of the read past the breakpoint in the bottom left.

**Supplemental Figure 5:**
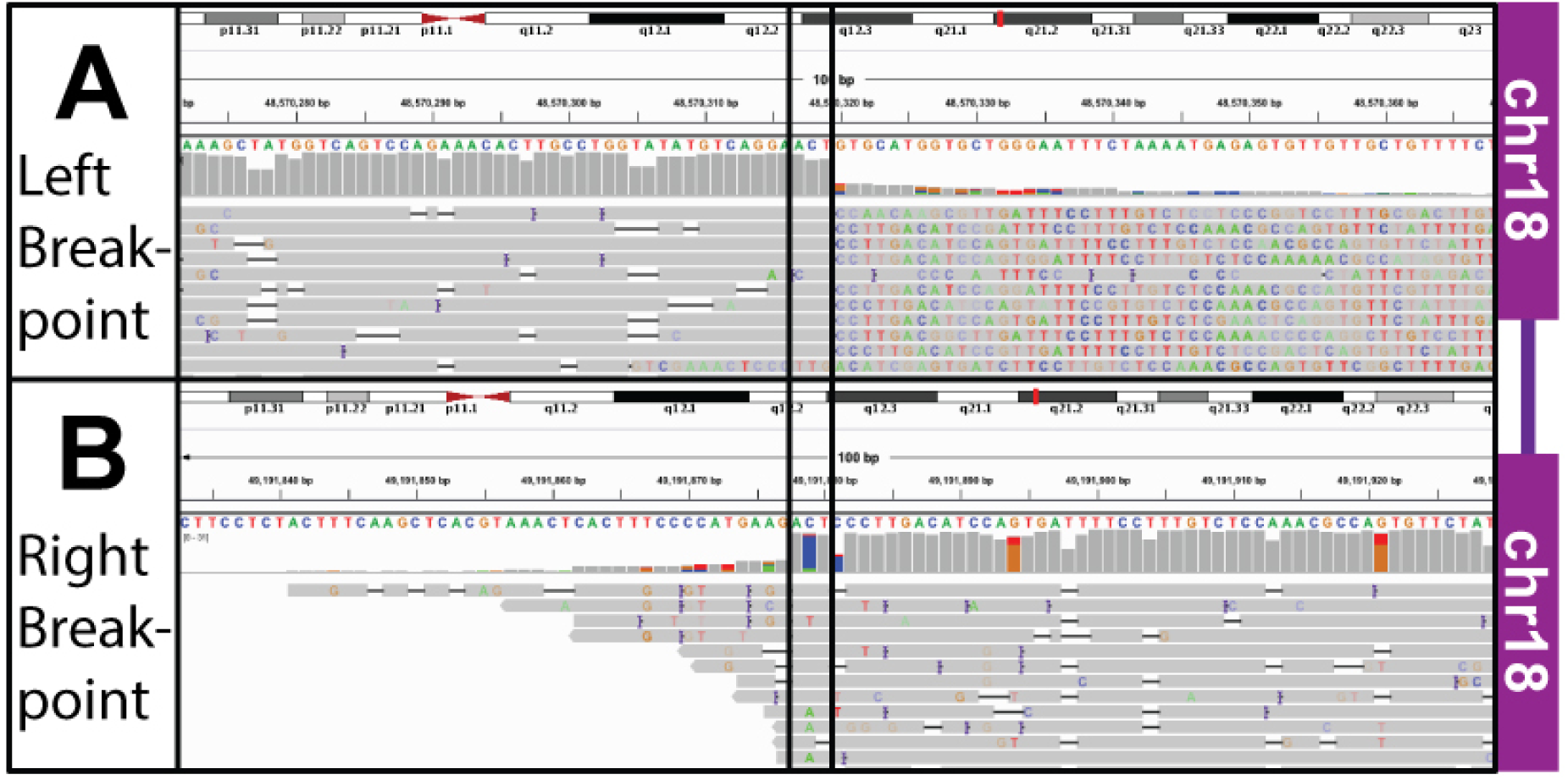
IGV Screenshot of Interstitial Deletion (SV05) alignment. The plot shows the alignment to the area upstream A) and downstream B) of the deletion in chr18. Note the erroneous extension of the read past the breakpoint in the top right.

**Supplemental Figure 6:**
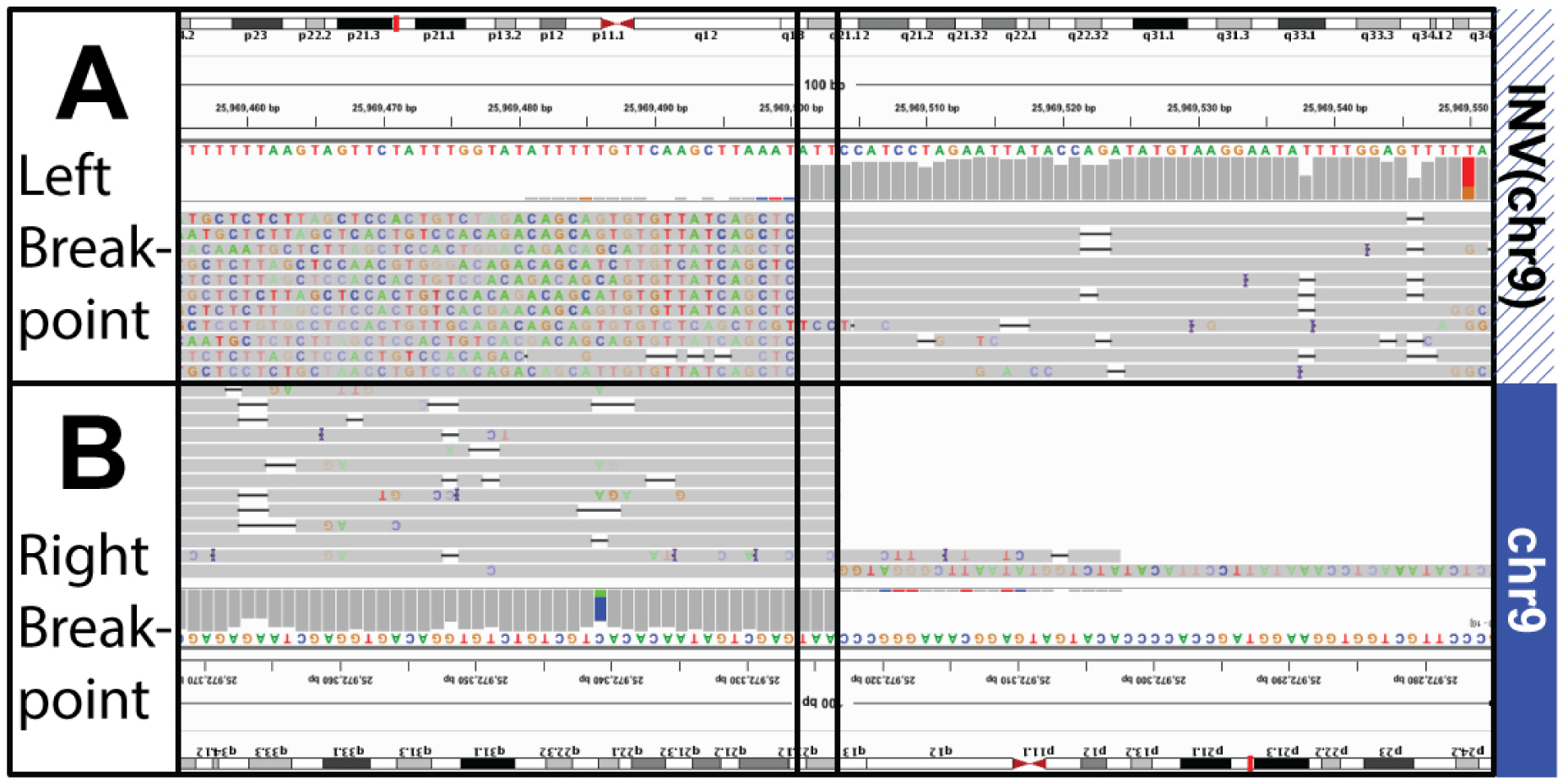
IGV Screenshot of Inversion (SV09) alignment. The plot shows the alignment to the inverted area A) and B) the area downstream of the inversion. We have flipped B) to show how the two parts align. Note the erroneous extension of the read past the breakpoint in the top left.

